# The calculation of the mutation frequency for humans at different proton doses in a mathematical model and estimation of the mutation risks for the space explorations

**DOI:** 10.1101/2025.01.28.635376

**Authors:** Metehan Kara, Melahat Bilge Demirköz

## Abstract

DNA is considered a fundamental component of life, yet it remains vulnerable to damage under extreme conditions, such as ionizing radiation exposure. To better understand this fragility, it becomes important to estimate mutation frequencies under different radiation doses. Furthermore, this approach has potential for future applications, especially in the context of deep space exploration, where astronauts are exposed to higher levels of cosmic radiation. For this purpose, we developed a mathematical model by integrating two existing models, the Monte Carlo Excision Repair (MCER) model and the Whack-a-Mole (WAM) model, both specifically adapted for use in manned space missions. The WAM model is supported with the Monte Carlo simulation to address the lack of human experimental data available in previous studies so that by calculating four key variables related to the human cells defined in the WAM model, potential mutations in astronauts during space exploration were estimated. The results showed small deviations from previous studies, which can be attributed to differences in the type of radiation sources as well as the organisms studied being different from those used in previous studies. With this study, researchers can now better predict mutation frequency during deep space missions by considering the impact of cosmic radiation. This is particularly important in the context of future missions to the Moon and Mars, where cosmic radiation will play an important role in mission planning and risk management.

## Introduction

Radiation exposure has left a deep trauma in human history as a result of some events and accidents in the nuclear age. For this reason, understanding the effects of radiation on living things has been a top priority for the scientific community since the beginning of the 20th century[1]. One of the landmark studies was Joseph Muller’s mutation experiments on the fruit fly between 1910-1915 at Columbia University, which led to the emergence of the WAM model. In Muller’s experiment, he observed the mutation effects caused by radiation on the eggs and sperms. After many successful cultures of Drosophila, he stated that these mutations could not arise from spontaneous phenomena, but from high-energetic radiation such as X-rays[2]. In addition to this experiment, another influential study played a significant role in the emergence of the WAM model[3]. A study called the “mega mouse” project conducted by Russell and Kelly at Oak Ridge National Laboratory at 1950s investigated radiation effects and changes in the genetics of mice. They worked on defining the set of loci to observe the amount of change intended to calculate the mutation rates in germ lines under radiation exposure. The procedure that was applied was relatively simple. The male mice population was divided into two groups. One group was affected by radiation such as X-rays whereas the other one was not subjected to any kind of radiation. Then, these two groups of males mated to the “T-stock females” so that one can observe the mutation occurred in any specific loci due to recessive mutations. With the help of this distinguishing aspect, Russell could measure the mutation rate at specific loci[4].

After these major efforts in molecular biology and the beginning of the space race, the scientific community began to investigate the limits of life in space and the effects of deep space travel on the lives of astronauts. First of all, there are three main sources of radiation that affect life in space: the Galactic Cosmic Ray (GCE) and the Solar Particle Event (SPE) and trapped radiation around the planets that have magnetic fields. The galactic cosmic ray is composed mostly of high-energy particles that come from outside the solar system and travel through the entire interstellar space. It consists of mostly protons (around 90%), helium and heavy particles. In addition, solar particle events (SPE) can be defined as high-energy particles emitted from the sun. As with the GCR, the majority of SPE is made up of protons (around 90%), with a certain proportion of helium and heavy particles. Although they share some common features with GCR, they have some differences in origin and energy spectrum[5]. Both Galactic Cosmic Rays (GCRs) and Solar Particle Events (SPEs) include high-energy particles, especially protons, which have the ability to ionize biological materials and thereby pose significant biological hazards[6]. This ionization can result in complex DNA damage, increased mutation frequencies and elevated risks of cancer, cardiovascular disease and neurological disorders in living organisms[7].

As a consequence, a comprehensive understanding of the effects of these radiation sources on biological systems is crucial for the effective planning and safety of space missions. Furthermore, understanding the biological effects of GCRs and SPEs can provide information about the design of life sustainment systems and habitat structures, helping astronauts to maintain optimal health during their long-term missions in space. By investigating multidisciplinary research involving radiation biology, materials science and aerospace engineering, mission officer can ensure safer and more resilient environments for human exploration of deep space missions to the Moon, Mars and beyond. Ultimately, a thorough understanding of how GCRs and SPEs affect living organisms is crucial for assuring a sustainable and long-term human presence in extraterrestrial environments. In the scope of space missions, ionizing radiation exposure from Galactic Cosmic Rays (GCRs) and Solar Particle Events (SPEs) can induce a variety of genetic mutations in biological organisms. Such mutations originate through the interaction of high-energy particles with cellular DNA, and their severity levels are significantly affected by the Linear Energy Transfer (LET) of the radiation. LET describes the amount of energy that a radiation particle transmits its energy to the material through passing per unit distance[8]. High LET radiation, such as the heavy particles found in GCRs, tend to induce dense ionization along their path, resulting in complex and clustered DNA damage. On the other hand, low LET radiation, typically associated with SPEs such as protons, causes more dispersive ionization events, resulting in simpler mutations such as single-base substitutions, small insertions or deletions [9] These mutations are generally much simpler to be repaired by cells, but they can still contribute to long-term health risks such as cancer and genetic instability. Moreover, high LET radiation can reduce cell viability by causing lethal mutations at a higher frequency, while low LET radiation can increase the likelihood of non-lethal mutations that accumulate over time [10]. It is critical to understand the relationship between LET and mutation types to assess the genetic risks faced by astronauts during long-duration missions to destinations such as the Moon or Mars.

Based on current knowledge, it is an effective initial approach to assess the biological risks associated with deep space missions to focus on proton-induced mutations since protons are a major component of space radiation, especially during Solar Particle Events (SPEs), and their interaction with biological tissues can lead to significant DNA damage, including single base substitutions and small insertions or deletions [9]. By concentrating research efforts on understanding how protons lead to genetic mutations, scientists can develop targeted protection strategies and medical countermeasures to protect the health of astronauts. This baseline work not only helps minimize immediate radiation hazards but also provides crucial information that advances the design and planning of long-duration missions to destinations such as the Moon and Mars. As the mission plans proceed, the extension of the studies to incorporate the effects of heavier ions will further enhance the effectiveness of protective measures and enable comprehensive safety for future deep space explorers.

### Whack-a Mole (WAM) model

The WAM model is a mathematical model to guide the researchers in calculating mutation frequencies of certain species such as mice or *Drosophila*. The crucial feature of this model is that it relies on dose rate essentially, as opposed to previous models that concentrate on calculating the mutation rate dependent on radiation dose, instead of dose rate. In this model, first of all, the system starts with definition of time dependent number of normal cells and mutated cells as described below[2].

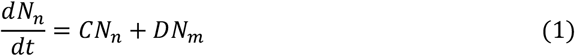

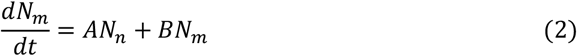

In the above equation, A, B, C, and D are fixed coefficients for different conditions. In here, the mutation is defined as the permanent unrepaired DNA damage in a DNA sequence so that there is no going back from mutated cells to normal cells[11]. In other words, the coefficient D that symbolizes as recovery is equal to zero(Wada et al., 2016). Also, t = 0 is defined as the initial moment when the radiation is absent from the environment, and at that time, 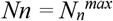.Also, the mutation frequency, F(t), is defined as 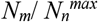.After that, considering the F(t) and the division of *N*_*n*_ in both sides of equation 2 leads to equation as below.

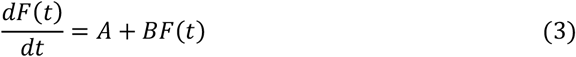

In the equation above, the symbol “A” represents the creation of mutated cells, whereas the “B” denotes the decay effects on the cell, and the A and B terms can be defined as below[11]:

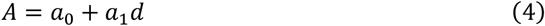

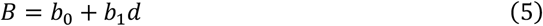

In here, “a_0_” stands for the spontaneous mutation effects; and “a_1_” mutation effects under radiated conditions; “b_0_” cell death effects under natural conditions; “b_1_” cell death effects under the radiated conditions and d represents constant dose rate. In here, since B terms are decreasing effects on mutated cells, B term’s sign should be changed to minus.

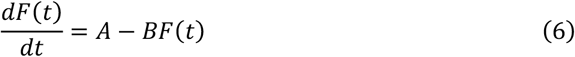

Then, by the applying the terms in the equation 6, the obtained formula is shown below.

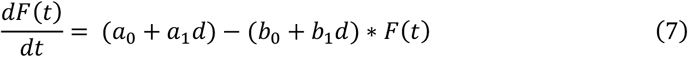

Moreover, the stationary condition is defined as below:

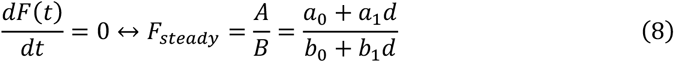

and consideration of equation 4,5 and 8, the analytical solution of equation 6 is below [2].

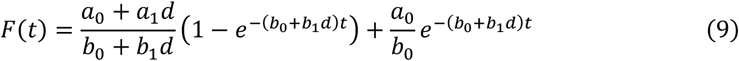

Also, above equation is widely used for describing plant and animal growth [11].

### Monte Carlo Excision Repair (MCER) model

Monte Carlo Excision Repair (MCER) model is a computer-based algorithm to allow researchers to determine the mutation rate after a specified number of cells, with particular energy level, under a certain radiation particle type and any dimethyl sulfoxide (DMSO) concentration. It relies on a well-known Monte Carlo simulation that describes the DNA damage under specific radiation exposure, but this model also defines the DNA repair mechanism and provides information about the DNA repair rate[13]. With the help of this information about the knowledge of DNA repair rate and its repair cycle, the DNA mutation rate can be obtained from the simulation output. Several DNA repair pathways in the MCER model give this mutation rate. For example, in MCER, there are short patch BER (base excision repair), long patch BER and Nucleotide excision repair (NER). The mutation rates can be examined by looking at the DNA repair pathways one by one and the mutation rates in different scenarios[14]. For instance, the scenario might be SP BER/ NER or LP BER/ NER so the mutation rates will be different from the DNA single repair pathway. Therefore, the MCER algorithm first forms the DNA damage at one of the DNA strands and then decides the DNA cluster so that computer codes can determine the average distance between damage sites of the DNA[15]. After that, the algorithm determines whether the repair pathways and patches are short or long. Then, base substitutions occur, and finally, the patch is complete, and the simulation will be terminated so that one can reach the outcomes of the previously determined radiation particle type and its energy. However, since this model relies solely on the base substitutions, unfortunately, it does not include insertion and deletion types of mutations[14]. In the future applications, these types of mutations can be easily integrated to the model.

## Material and methods

First, we adjusted the MCER simulation which includes 10,000 mammalian cells and the radiation particle type as a proton, due to its common presence in the space radiation environment. Moreover, the energy ranges of MCER are chosen between of 0.2 MeV to 5.5 MeV to investigate immediate biological impacts of space radiation, and the DMSO concentration is set to 0 to prevent any scavenging effects on radical components caused by the radiation. We did not only take the SP BER repair pathway to remove any additional mutation generated by nature of the repair scenarios, but also took the results caused from the damage type as “all”. After obtaining all data points from the MCER simulations, we have mutation rates in humans according to the SP BER and damage type as “all”. However, in MCER simulation results, the results, which can be DNA damage or mutation rate, are obtained from the particle energy not from the radiation dose or dose rate[14]. Therefore, since in the WAM model, the formula depends on radiation dose rates [2] we needed to convert all data points’ energy levels into the dose and its rates. For this purpose, we assigned the DNA components, their size and density to the software called SRIM-TRIM software [16]. Hence, with the help of the SRIM-TRIM software, we can convert all the proton energy from 0.2 MeV to 5.5 MeV of data points obtained from the MCER simulation to radiation dose rates in order to compare against the WAM model. In order to fit the WAM model to the data, we used the least squares fit in R language. Then, according to the four variables obtained as a result of these processes, the mutation frequency that will occur in astronauts with different scenarios that may occur during space travel was calculated.

## Results

In this study, we combined the MCER simulation and WAM model in the literature with R codes in order to find the mutation frequency that will develop in humans in a radiation environment. According to the study, the average cosmic radiation dose rate on a Mars journey is 1.85*10^−5^ Gy/h [17]. As an example, based on this radiation dose rate, the fit of WAM model to the simulation output of MCER and the number of mutations that will occur in 10,000 mammalian cells of astronauts during a Mars journey were given below.

**Table 1.** The mutation frequency that occurs in 10,000 mammalian cells of a human with the proton radiation at 1.85*10^−5^ Gy/h radiation dose rate.

| Total Absorbed dose<br>(Gy) | Mutation<br>Frequency | Total Absorbed dose<br>(Gy) | Mutation<br>Frequency |
| --- | --- | --- | --- |
| 4.498 | 5.02E-06 | 0.192 | 2.17E-06 |
| 3.885 | 4.79E-06 | 0.168 | 1.97E-06 |
| 3.453 | 4.59E-06 | 0.146 | 1.89E-06 |
| 2.906 | 4.40E-06 | 0.126 | 1.82E-06 |
| 2.460 | 4.23E-06 | 0.110 | 1.76E-06 |
| 2.066 | 4.06E-06 | 0.0968 | 1.71E-06 |
| 1.618 | 3.80E-06 | 0.0809 | 1.66E-06 |
| 1.293 | 3.55E-06 | 0.0806 | 1.60E-06 |
| 1.056 | 3.34E-06 | 0.0481 | 1.56E-06 |
| 0.888 | 3.23E-06 | 0.0380 | 1.50E-06 |
| 0.737 | 3.01E-06 | 0.0304 | 1.403E-06 |
| 0.583 | 2.88E-06 | 0.0248 | 1.37E-06 |
| 0.438 | 2.65E-06 | 0.0204 | 1.32E-06 |
| 0.336 | 2.45E-06 | 0.0170 | 1.30E-06 |
| 0.209 | 2.30E-06 | 0.0143 | 1.26E-06 |

**Figure 1.**
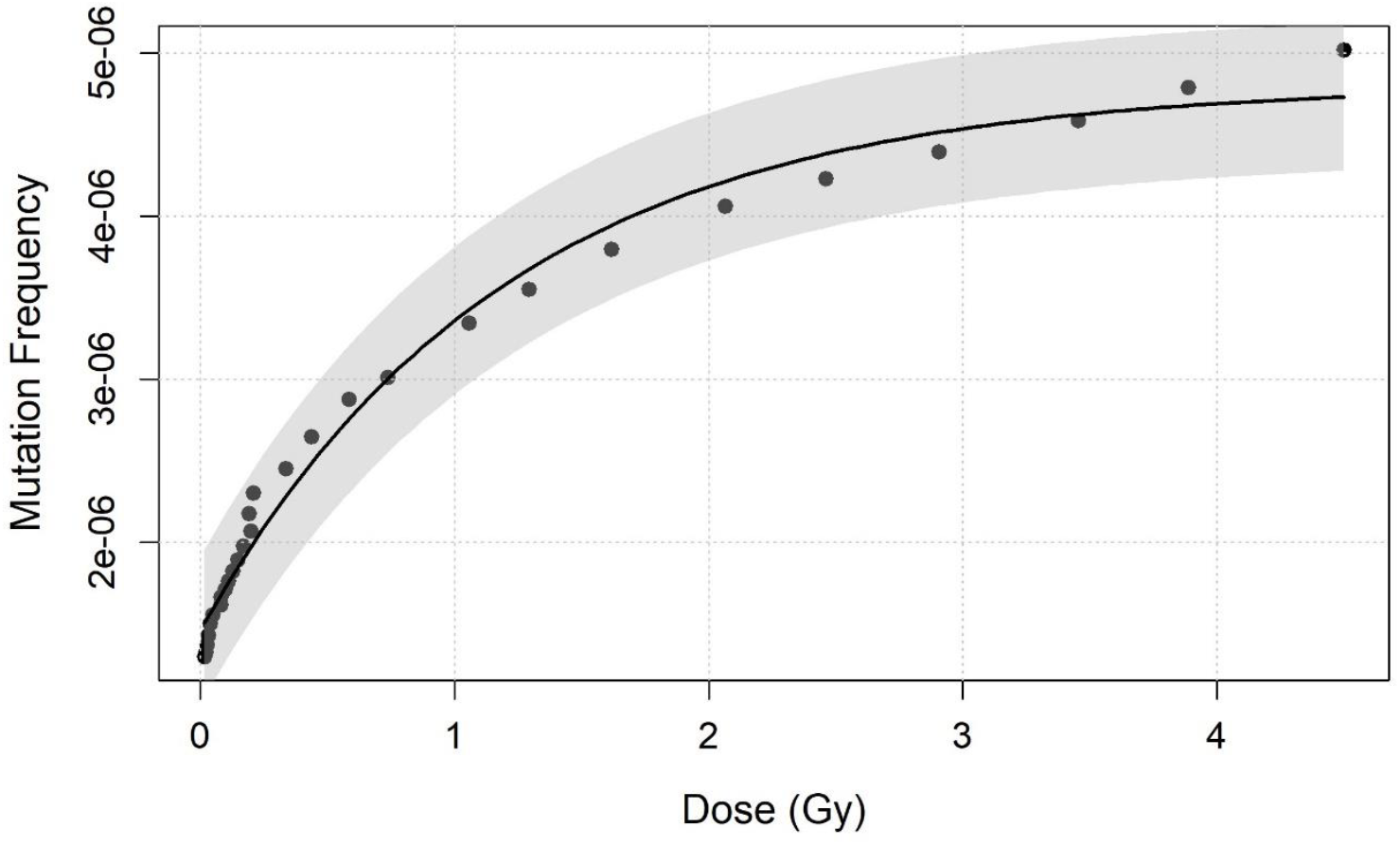
The simulated mutation frequency from MCER against the absorbed dose for a proton irradiation with dose rates as 1.85*10^−5^ Gy/h, fitted with WAM model curve from the MCER simulation with 95% confidence interval shaded area around curve.

**Figure 2.**
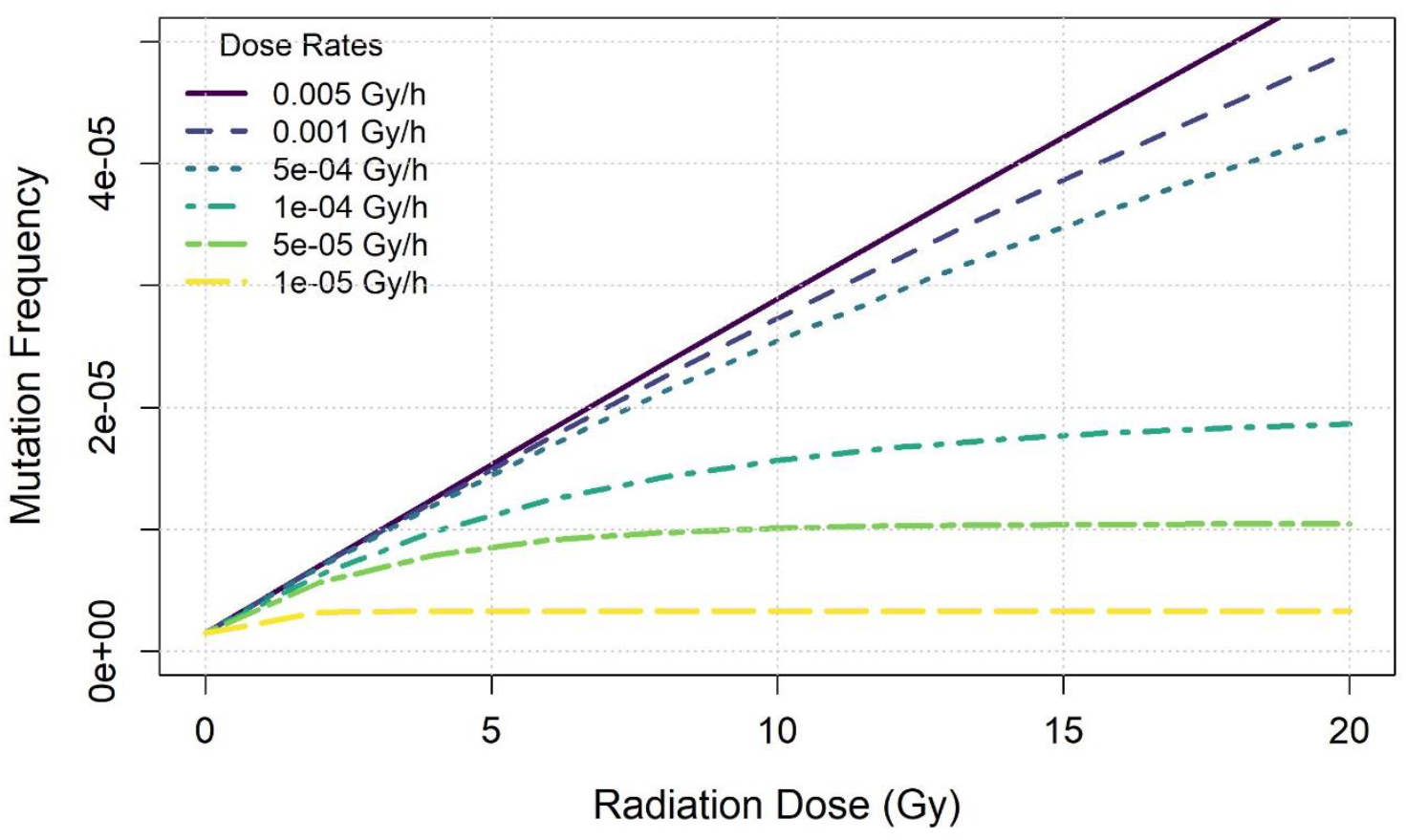
Representation the mutation frequency defined in a vertical axis with different radiation dose levels in horizontal axis according to the different radiation dose rates

**Figure 3:**
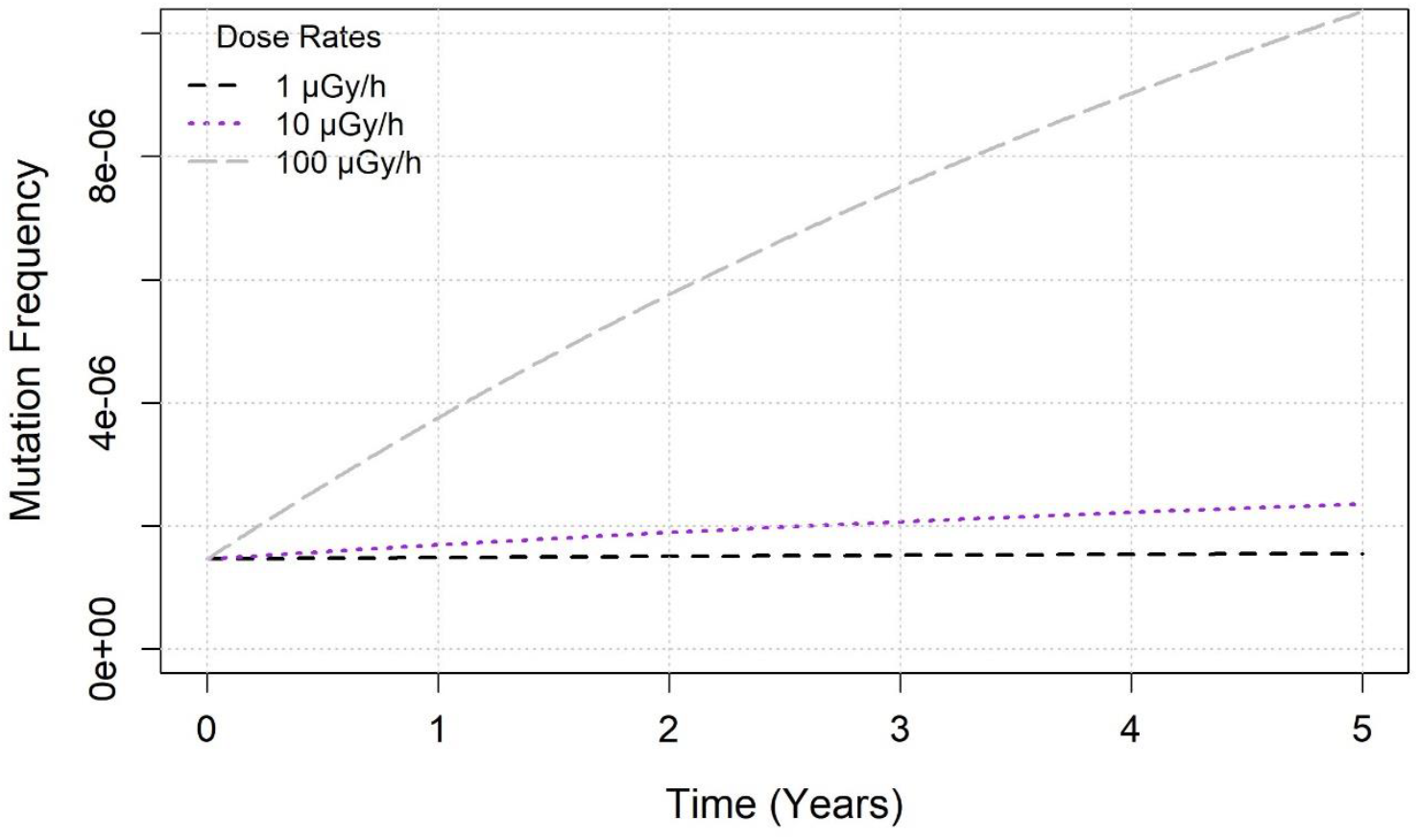
Representation the mutation frequency defined in a vertical axis with different times in horizontal axis according to the different radiation dose rates

**Figure 4.**
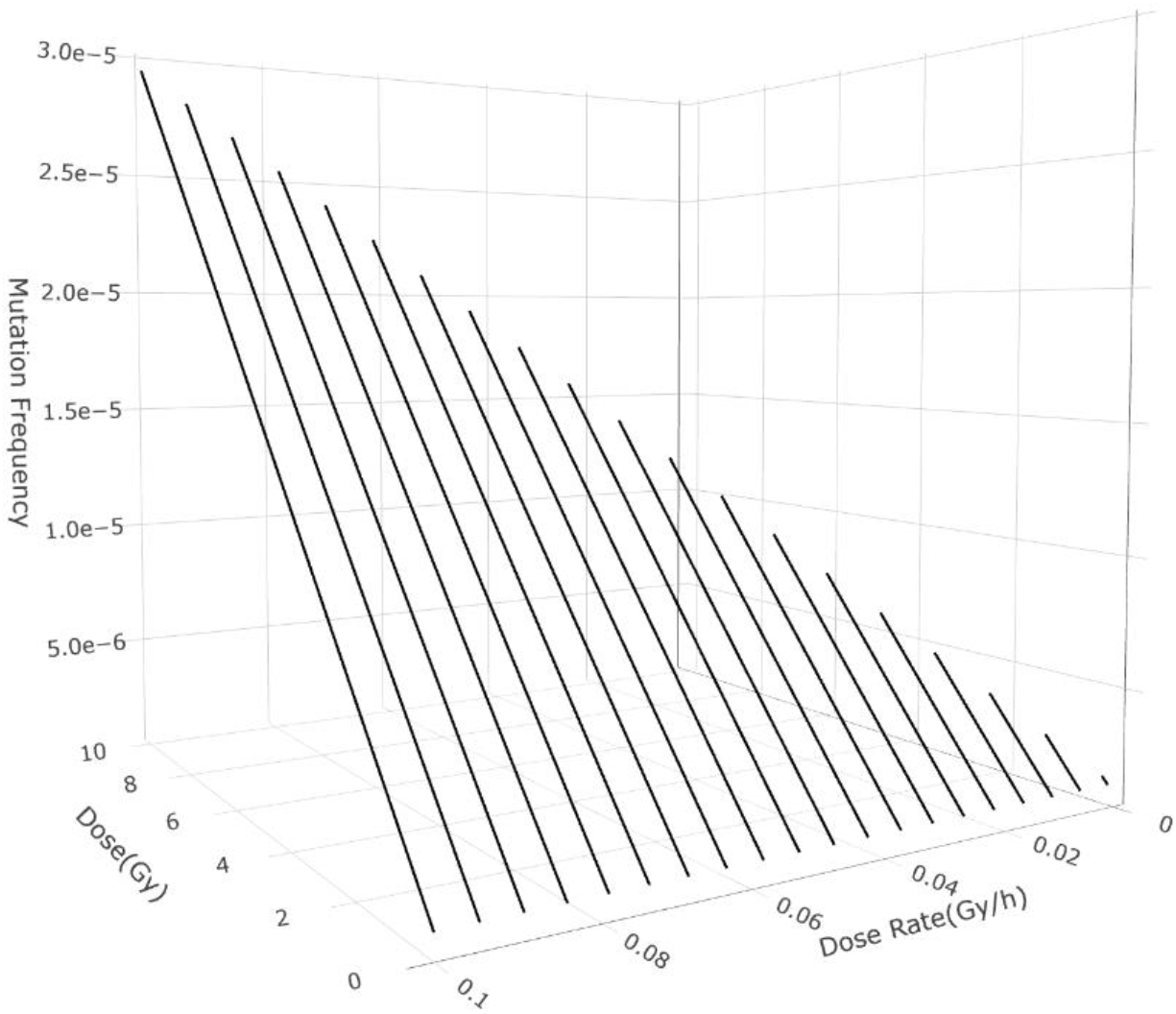
3D representation the mutation frequency with different radiation dose and dose rate

After all these fitting procedures, we obtained four coefficients for human at 1.85*10^−5^ Gy/h dose rate as below:

**Table 2.** The value of four coefficients based on 1.85*10^−5^ Gy/h dose rate.

| Coefficient | Value |
| --- | --- |
| $a_0$ | $2.25\text{E-}11$ [/hour] |
| $a_1$ | $2.85\text{E-}06$ [/Gy] |
| $b_0$ | $1.24\text{E-}05$ [/hour] |
| $b_1$ | $1.91\text{E-}03$ [/Gy] |

Indeed, these coefficients were found from a 1.85*10^−5^ Gy/h dose rate. Hence, they will change slightly under different low dose rates.

After finding these coefficients values, our mathematical model can be compared with literature data either human cells or mouse cells with different proton energy range by considering the dose rate of the data samples [18,19].

In addition to comparison of these samples with our mathematical model in a very short time, the comparison between the samples and our mathematical model in long time period can be seen as below [20,21].

In addition to these comparisons, according to the already founded coefficients value, the mutation frequency that may occur in some possible human space exploration scenarios is shown as below.

## Discussion

After all this fitting procedure that combined MCER data points into the WAM model’ mathematical formulation, we intended to obtain the four coefficients necessarily required to calculate the mutation frequency especially for humans because when one can examine the WAM model and its outcome for the coefficients and mutation rates, there is only specific species data [2,3]. The researchers that establish this WAM model benefited from the previous experiments that define radiation dose rate and related mutation rates. However, since the radiation experiments that are conducted on whole body living creatures, especially humans, can be ethically problematic, there is no excessive data obtained from humans in this version of the WAM model. For this reason, we aimed to develop the WAM model further to be able to make predictions for humans. As mentioned above, there is no such excessive experimental data about human mutation rates under different radiation dose rates. Therefore, we supported the WAM model formula with the MCER simulation data that include the human mutation rates according to the given dose rates [14]. We obtained human mutation rates data that fitted to the WAM model’s mathematical formulation. After that, with the help of the R fit codes, we connected these two elements. Our coefficients are different from the traditional WAM model as expected; since the mutation frequency is given from the consideration of DNA structure not from the whole organismal level. That is, the mutations in MCER are defined as the replacement of the bases between DNA strands after radiation exposure[14]. On the other hand, in the WAM model, the researchers utilized experiments that included the mutation effects on *Drosophila* and mouse during the radiation as a biological organism [11].

In addition, one of the reasons why our data is different from the WAM model is due to what our radiation type is because in this study we selected the radiation type as a proton since it is the dominant component of the penetrating radiation in the space environment[22]. However, the WAM model exploited several radiation types such as X-rays, neutrons or gamma rays [2]. While the biological outcome can be the similar, there are differences in damage mechanisms [23]. That is, ionizing radiation can affect the organisms diversely according to the ionized radiation types [24]. Moreover, if the differences between X-rays and protons are investigated, there are serious characteristic differences to between[25]. For instance, an X-ray or a gamma ray can be thought as a bullet that may pass through the body, with some probability that it stops, imparting all its energy in the body. The proton is a heavy and positively charged particle that loses energy along its path and travel shorter or can penetrate further than an X-ray, depending on its energy. Close to the end of its path inside the material, it can lose its energy catastrophically as defined by the Bethe-Bloch formula, in the so-called the “Bragg peak”[26]. Therefore, proton radiation can cause more damage by creating DNA strand breaks and cytotoxicity [27].

Considering these, it is expected that the coefficients obtained in this study would differ from those in the WAM model. Therefore, upon investigating the first table, it can be seen that the mutation frequency increase as the radiation dose rises. From lowest radiation dose to highest dose, the mutation frequency differences are around 4-fold. Then, analyzing the upper limit of radiation and its mutation frequency in this study, it becomes evident that a high mutation frequency was observed, which is reasonable based on the literature because human cell death will be anticipated at such radiation dose and mutation frequency [28]. Upon reviewing the first figure, it can be seen that the fitting procedure between the WAM model and MCER simulations suits each other within the confidence interval. Furthermore, when the second figure is examined, the mutation frequency is proportional to the radiation doses and dose rates. While the radiation dose increases, the highest dose rate trajectories become linear relation while the lowest ones are more saturated curves around the highest radiation dose. These patterns are highly similar to previous study [11], that is logical consideration due to similarity between human and mouse cells. Furthermore, when we look at third and fourth figures, we observe that the mutation frequency increases as the radiation dose rate rises with based on time and radiation dose, but it is evident that this rate is much lower especially when the lower dose rates are observed. This may indicate that organisms have the capacity to repair more efficiently at low radiation doses as suggested by research in the past [29]. These findings are also correlated with the previous study [11].

Moreover, after finding the coefficients value by fitting of simulation results to WAM model, this new model belonging to the human cells can be compared to the data already conducted in the literature. However, unfortunately, there is no extensive data information in the literature available that contains proton radiation dose, dose rate, time and mutation frequency. Nevertheless, there are several experimental results that contain proton radiation dose/dose rate or space condition, mutation frequencies (TK mutants or lacZ mutants) and time [20,21]. The figure 5 and 6 describe the comparison of our model with experimental data. In here, the most significant data point is the one in figure 6, which shows the 134-day “Kibo” mission that was conducted on the ISS during the years 2008 and 2009[20] It is clear from this data that our model provides a suitable model for explaining this experiment. However, a single study is not enough to demonstrate the accuracy of our model. For this reason, we compared our model in both data sets with short time intervals and relatively long experimental results, similar to the “Kibo” mission. In both figures, it is shown that the experimental results are in good agreement with the obtained model. Although the curve obtained only in figure 5 is thought to show a straight pattern, which is reasonable given the very short time interval. Nevertheless, this model seems to be more effective at lower dose rates and longer time intervals.

**Figure 5.**
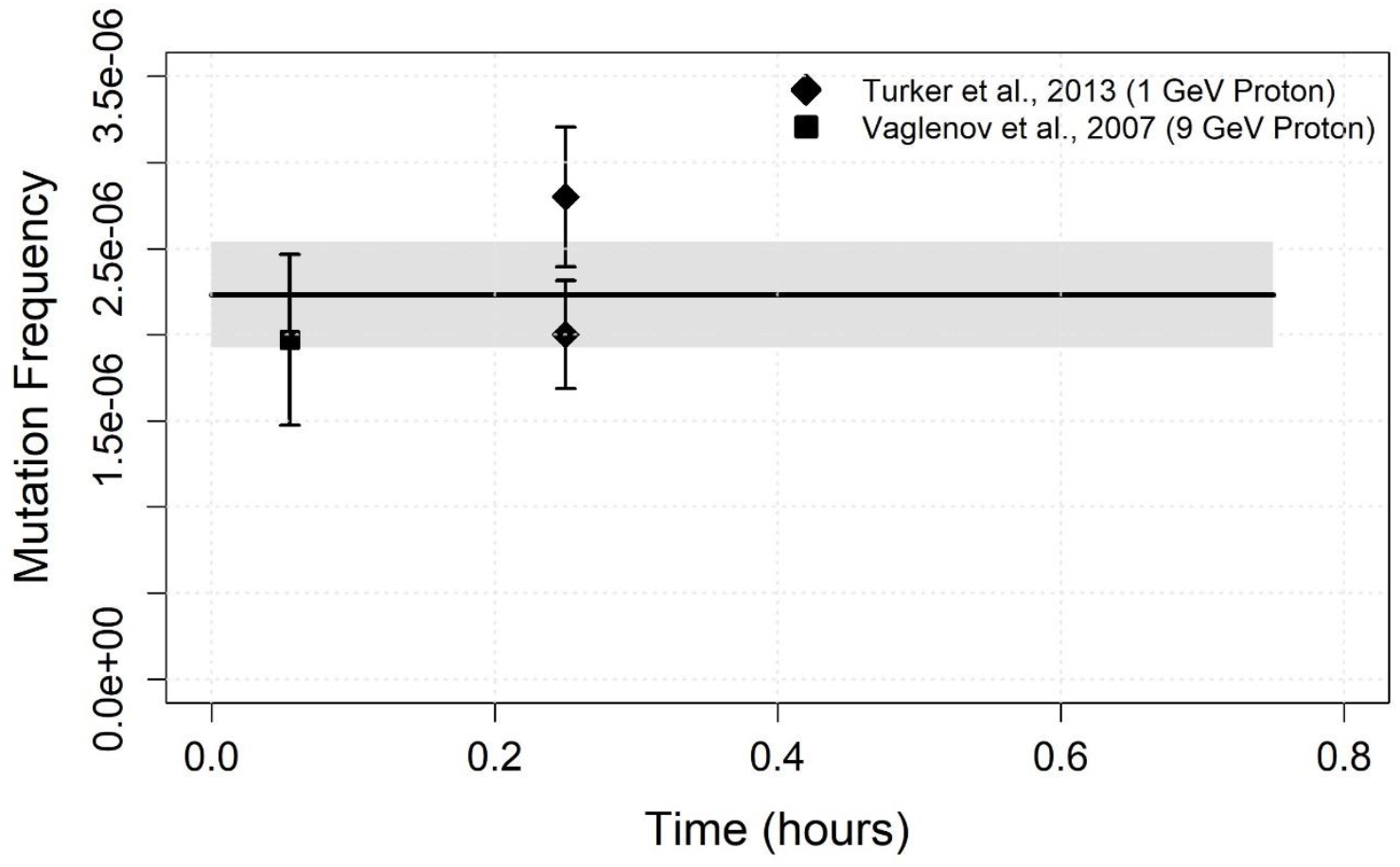
Comparison of mutation frequency of different samples in very short time period.

**Figure 6.**
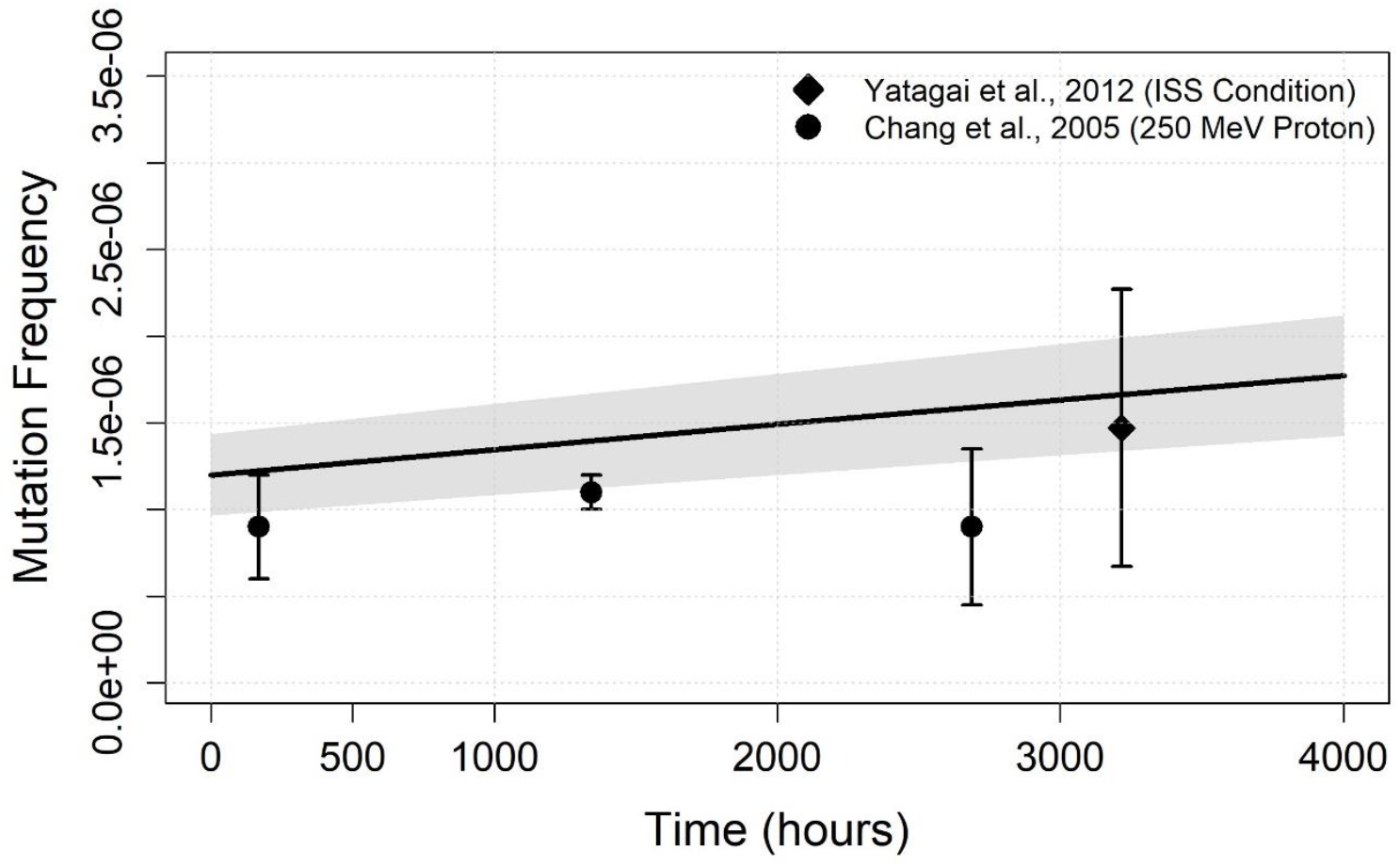
Comparison of mutation frequency of different samples in long time period

After comparing our model with experimental data reported in the literature, we assessed the mutation frequencies that may occur during potential space missions due to cosmic radiation exposure. The results, presented in Figure 7, demonstrate a linear increase in mutation frequency over time. Notably, for a Mars mission, the estimated mutation frequency is 2 to 2.5 times higher than the initial condition. Therefore, it is better to reduce either the mission duration or the radiation dose and dose rate received by astronauts to ensure the health and success of the mission.

**Figure 7:**
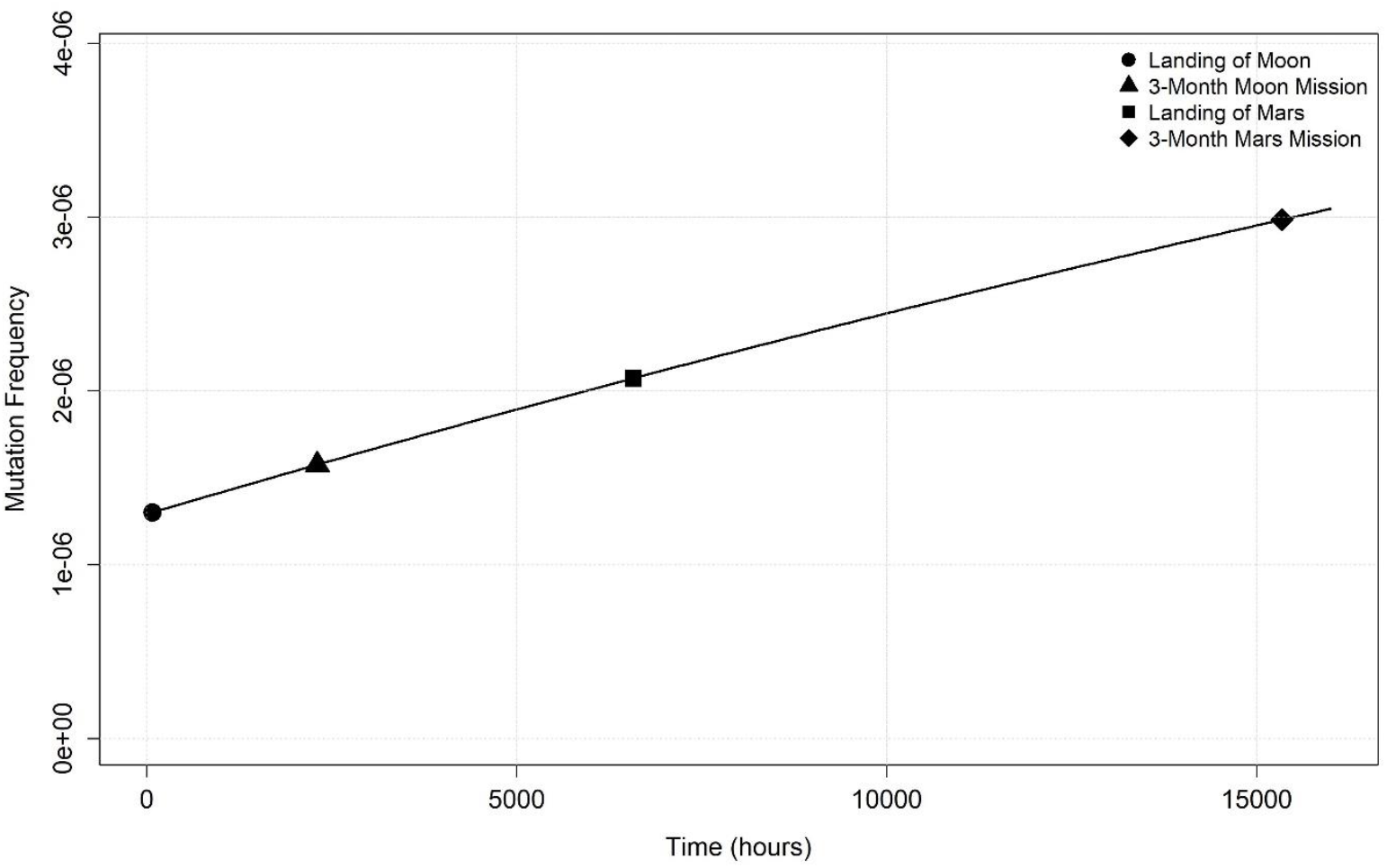
The estimation of mutation frequencies in possible space missions over time

Overall, the MCER simulation reveals the mutation frequency that occur in humans in a radiated environment. Since, the MCER simulation provides the mutation rate as a chemical event -base substitutions- and the radiation type that we chose in this paper may have different effects on DNA and living things, unlike the WAM model. Therefore, considering these key aspects, the results that we obtained in this study can be different from the WAM model based on sufficient reasons. Also, planning space program research with proton-induced mutations is a potentially useful way to approach it because protons are the most common component of space radiation, especially during Solar Particle Events, which pose significant genetic risks to astronauts [6]. This targeted focus provides a strong groundwork for addressing more complex types of radiation in future deep space missions, enabling early development of effective shields and medical countermeasures.

As a conclusion, in this study, we developed a mathematical model to describe the generated mutation frequency for humans under radiation exposure by exploiting two models called WAM and MCER model. For the numerical results, all these coefficients named as a_0_, a_1_, b_0_ and b_1_ were found from the 1.85*10^−5^ Gy/h dose rate. That is why in a future study the researchers can utilize the technique that we described in this current study to apply to different scenarios such as calculating the mutation rate for the astronaut in a deep space mission or calculating the mutation rate of the local people under unwanted radiation exposure by adjusting the radiation dose rate and time [10,29].

